## Supplemental Data for "Glioma-derived M-CSF and IL-34 license M-MDSCs to suppress CD8^+^ T cells in a NOS-dependent manner"

**Table S1.** Antibodies used for flow cytometry. Marker, fluorophore, company, catalog number and dilution factors are reported.

| **Cell Marker:** | **Fluorophore:** | **Company:** | **Catalog #:** | **Dilution Factor:** |
|---|---|---|---|---|
| CD3 | BV510 | BioLegend | 100234 | 1:100 |
| CD4 | FITC | BioLegend | 100510 | 1:100 |
| CD8 | PE | BioLegend | 100712 | 1:100 |
| CD45 | Alexa700 | BioLegend | 103128 | 1:200 |
| CD11b | BV421 | BioLegend | 101251 | 1:100 |
| LY6C | BV785 | BioLegend | 128041 | 1:100 |
| LY6G | PerCP | BioLegend | 127654 | 1:100 |
| CD39 | PE | BioLegend | 143804 | 1:100 |
| CD73 | APC | BioLegend | 127210 | 1:100 |
| iNOS | APC | Invitrogen | 17-5920-82 | 1:100 |
| Viability Dye | Pacific Blue | Invitrogen | L34963 | 1:1000 |
| FcR Block | - | BioLegend | 101320 | 2 uL per sample |

**Table S2.** Cytokines, antibodies, and small molecule compounds used for expansion and inhibition of MDSC differentiation or inhibition of M-MDSC suppression. Company and catalog numbers are reported.

| **Cytokine/Antibody/Small Molecule** | **Company:** | **Catalog #:** |
|---|---|---|
| GM-CSF | R and D Systems | 415-ML-020 |
| M-CSF | R and D Systems | 416-ML-010 |
| G-CSF | R and D Systems | 414-CS-005 |
| TGF- β | R and D Systems | 7666-MB-005 |
| IFNγ | R and D Systems | 485-MI-100 |
| IL-1β | R and D Systems | 401-ML-005 |
| IL-2 | R and D Systems | 402-ML-020 |
| IL-6 | R and D Systems | 406-ML-025 |
| IL-12p40 | R and D Systems | 9807-IL-050 |
| IL-13 | R and D Systems | 413-ML-005 |
| LIF | R and D Systems | 8878-LF-025 |
| αIL-34 mAB | R and D Systems | MAB5195 |
| αM-CSF pAB | R and D Systems | AF416-SP |
| αPD-L1 mAB | R and D Systems | MAB90783 |
| αIFN-γ R2 mAB | R and D Systems | MAB773 |
| Apocynin | R and D Systems | 4663 |
| CGS 15943 | R and D Systems | 1699 |
| Celecoxib | R and D Systems | 3786 |
| Pexidartinib | R and D Systems | 7590 |
| Nor NOHA monoacetate | R and D Systems | 6370 |
| L-NMMA acetate | R and D Systems | 0771 |
| Methyl-D-tryptophan | R and D Systems | 5698 |

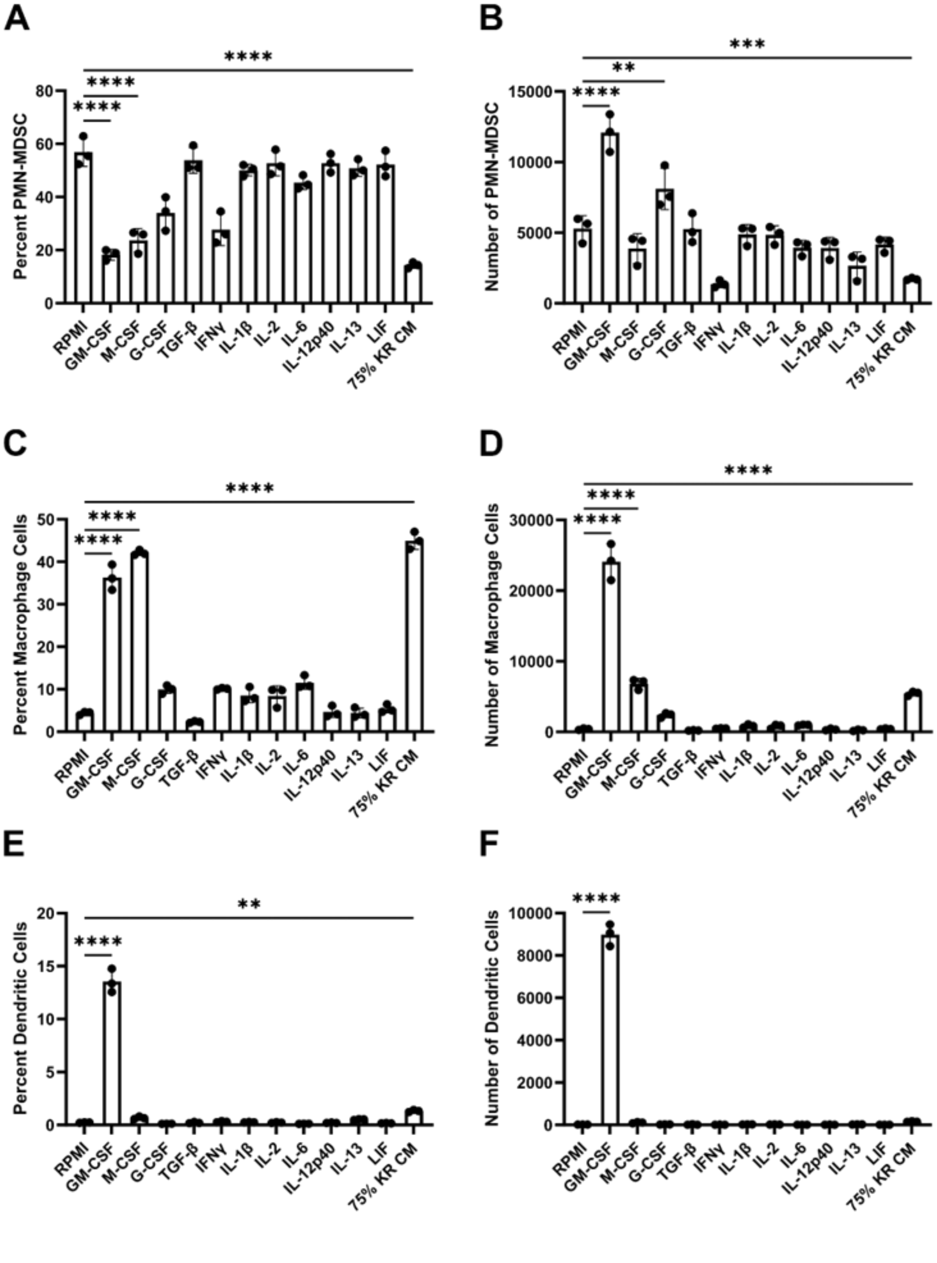


**Figure S1.** Glioma upregulated cytokines alter bone marrow differentiation ex vivo. Graph depicting the (**A**) percent and (**B**) numbers of Ly6C/Ly6G PMN-MDSCs generated from whole bone marrow cultured for 3 days in the presence of 40ng/mL cytokines or 75% KR158B conditioned media (n=3). Graph depicting the (**C**) percent and (**D**) numbers of F4/80 macrophages generated from whole bone marrow cultured for 3 days in the presence of 40ng/mL cytokines or 75% KR158B conditioned media (n=3). Graph depicting the (**E**) percent and (**F**) numbers of CD11c dendritic cells generated from whole bone marrow cultured for 3 days in the presence of 40ng/mL cytokines or 75% KR158B conditioned media (n=3). One-way ANOVA statistical analysis was conducted (Dunnett's multiple comparisons test). Differences are compared to the control condition or between cell lines. p-values: 0.0332(*), 0.0021(**), 0.0002(***)

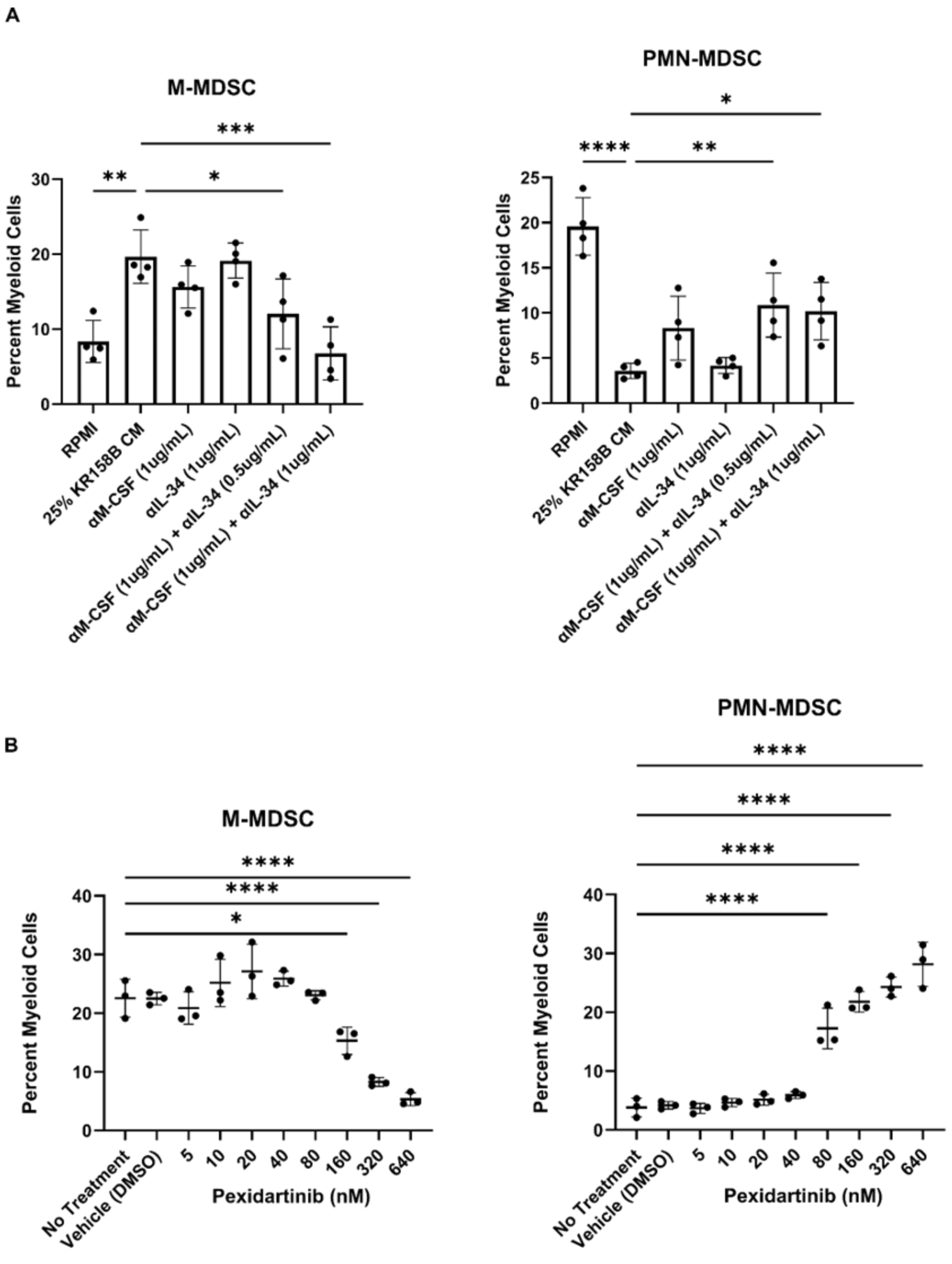


**Figure S2.** Inhibition of the CSF1R axis impacts percentage of M-MDSCs. (**A**) Whole bone marrow from wildtype mice were cultured in the presence of KR158B glioma conditioned media with M-CSF and IL-34 neutralizing antibody. M-MDSC percentage decreased to control levels with combination neutralization. PMN percentages were reduced in all conditions. (**B**) Whole bone marrow from wildtype mice was cultured in the presence of KR158B glioma conditioned media and the potent small molecule CSF1R inhibitor Pexidartinib. CSF1R inhibition resulted in a dose dependent decrease in M-MDSCs and an apparent increase in PMN-MDSCs percentage. One-way ANOVA statistical analysis was conducted (Dunnett's multiple comparisons test). Differences are compared to the control condition or between cell lines. p-values: 0.0332(*), 0.0021(**), 0.0002(***).

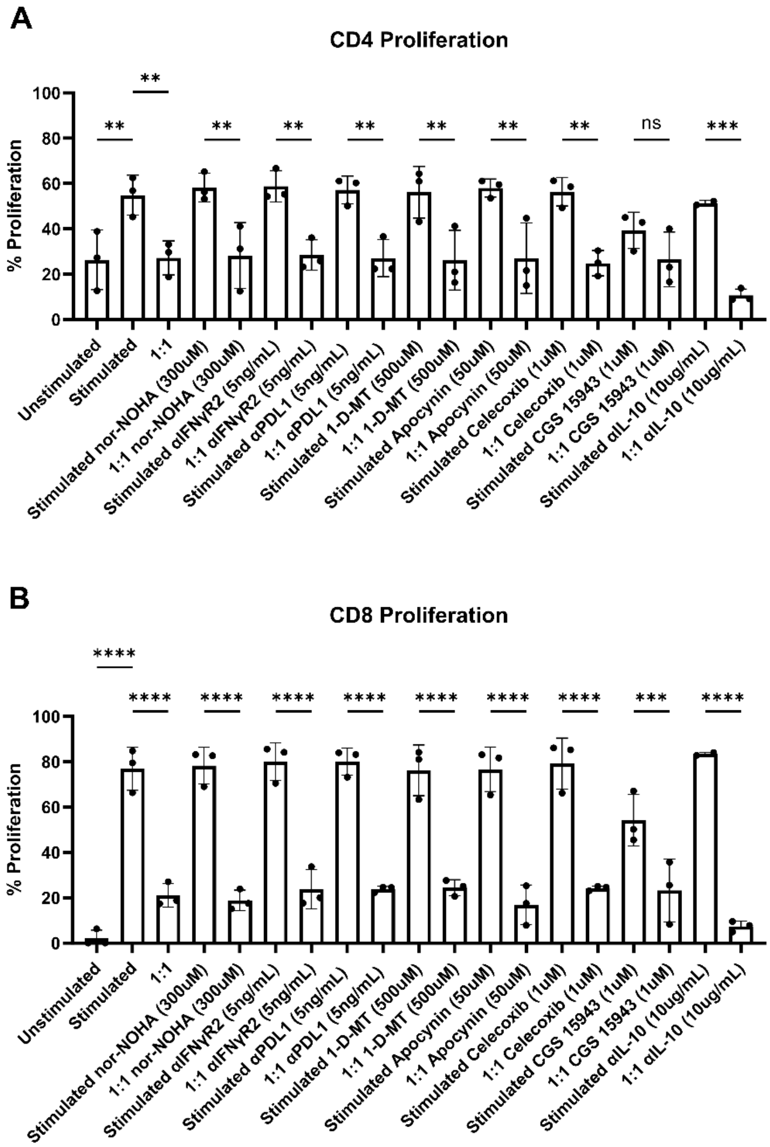


**Figure S3.** Inhibitor screen to identify mechanisms of glioma induced M-MDSC mediated T cell suppression. (**A**) Quantification of CD4 T cell proliferation in the presence of 1:1 ratio of M-MDSCs generated from KR158B conditioned media. Inhibitors targeting ARG1, IFNγR2, PDL1, IDO, NOX, COX2, A2AR, and IL-10 were tested to recover suppression induced by M-MDSCs. (**B**) Quantification of CD8 T cell proliferation. The tested inhibitors could not recover the proliferation of CD4 and CD8 T cells. One-way ANOVA statistical analysis was conducted (Dunnett's multiple comparisons test). Differences are compared to the stimulated control condition. p-values: 0.0332(*), 0.0021(**).

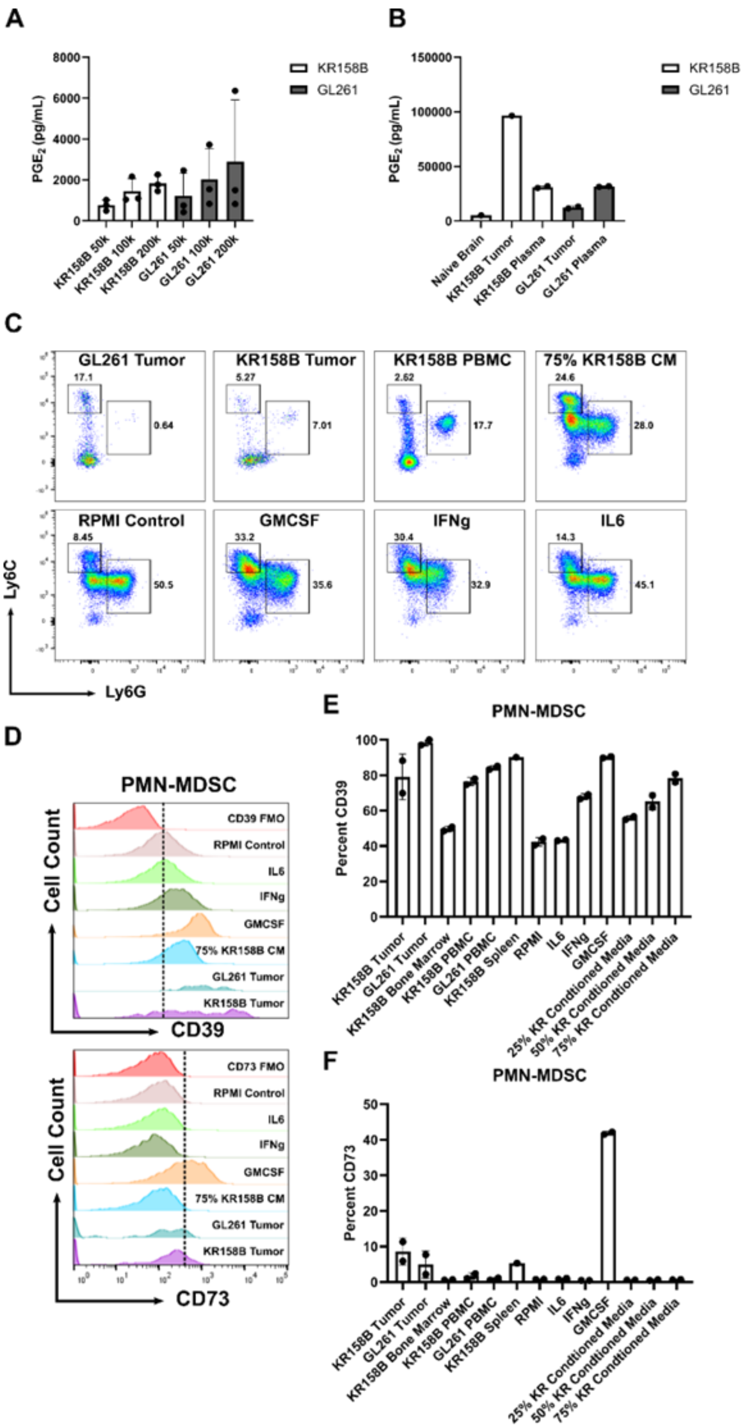


**Figure S4**. Glioma cells secrete PGE2 and PMN-MDSCs regulate CD39. (**A**) PGE2 ELISA on cultured GL261 or KR158B glioma conditioned media or tumors. (**B**) Representative flow cytometry analysis depicting gating strategy and percentage of M-MDSCs evaluated for CD39 and CD73 expression. (**C**) Representative flow cytometry histograms. (**E-F**) Graphs showing percentage of PMN-MDSCs expressing CD39 or CD73 on cell surface.

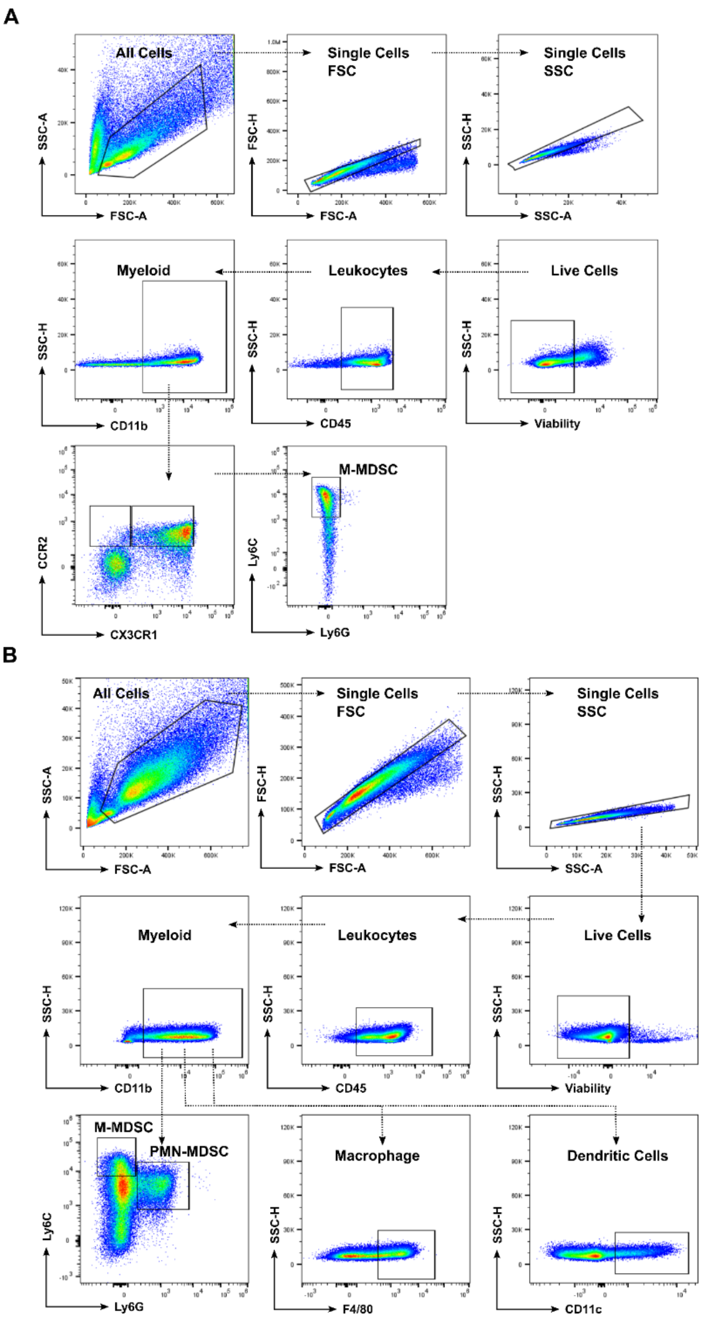


**Figure S5.** Representative flow cytometry gating strategy. (**A**) Flow cytometry gating strategy for CCR2/CX3CR1 expressing cells and CCR2/CX3CR1 expressing M-MDSCs generated from whole bone marrow cultured with KR158 conditioned media. Representative plot shows bone marrow treated with 75% KR158B conditioned media (**B**) Flow cytometry gating strategy for MDSC subsets, macrophages, and dendritic cells generated from whole bone marrow cultured with exogenous cytokines and KR158B conditioned media. Representative plot shows bone marrow treated with exogenous GM-CSF (40ng/mL).
